# Isolation of persisters enabled by ß-lactam-induced filamentation reveals their single-cell awakening characteristics

**DOI:** 10.1101/600700

**Authors:** Etthel M. Windels, Zacchari Ben Meriem, Taiyeb Zahir, Kevin J. Verstrepen, Pascal Hersen, Bram Van den Bergh, Jan Michiels

## Abstract

When exposed to lethal doses of antibiotics, bacterial populations are most often not completely eradicated. A small number of phenotypic variants, defined as ‘persisters’, are refractory to antibiotics and survive treatment. Despite their involvement in relapsing infections caused by major pathogens, processes determining phenotypic switches from and to the persister state largely remain elusive. This is mainly due to the low frequency of persisters in a population and the lack of reliable persistence markers, both hampering studies of persistence at the single-cell level. Problematically, existing methods to enrich for persisters result in samples with very low persister densities and/or a too high abundance of other cell types. Here we present a novel and highly effective persister isolation method involving cephalexin, an antibiotic that induces extensive filamentation of susceptible cells. We show that antibiotic-tolerant cells can easily be separated by size after a short cephalexin treatment, and that the isolated cells are genuine persisters. We used our isolation method to monitor persister outgrowth at the single-cell level in a microfluidic device, thereby conclusively demonstrating that awakening is a stochastic phenomenon. We anticipate that our novel approach can have far-reaching consequences in the persistence field, by allowing single-cell studies at a much higher throughput than previously reported.

## Introduction

Persisters are phenotypically distinct variants in a microbial population that survive a lethal antibiotic dose and are able to regrow after treatment ceases [1,2]. Given this population heterogeneity, interrogation of the persister physiology should rely on single-cell studies to properly capture their defining traits. However, these studies require considerable and fast enrichment of persisters as they are usually present at low frequencies and known to be in a metastable state. Problematically, apart from their antibiotic tolerance, no reliable marker currently exists to distinguish persisters from normal, susceptible cells. The state-of-the-art method to enrich for persisters involves lysis of susceptible cells by ampicillin, followed by sedimentation of intact persister cells [3]. Due to the poor separation efficiency during sedimentation, this method fails to efficiently remove cell debris and results in a persister density that is most often too low for microscopic studies. Furthermore, prolonged exposure of the culture to antibiotics or dead cell material could potentially affect persister formation [4–6]. The latter problem was addressed by Cañas-Duarte et al., who optimized a method to rapidly lyse susceptible cells using a chemo-enzymatic lysis solution [7]. Problematically, they did not validate antibiotic tolerance of their isolated cells, nor did they report the purity and density of the resulting sample. Other approaches using GFP expression, RpoS::mCherry expression, or the RNA-binding Thioflavin T as fluorescent markers for persistence, make too strong assumptions on the physiological state of persisters and therefore generate samples that are highly contaminated with normal, susceptible cells [8–10]. Attempts to enrich persisters using chemical pretreatment [6] or strains that are engineered to accumulate toxins [11] potentially generate artefacts that confound insights in naturally occurring persistence.

In this study, we established a novel, highly efficient persister isolation method that largely addresses the challenges posed by single-cell persistence studies. We show that persisters can be effectively isolated by filtration after ß-lactam-induced filamentation. Cells isolated in this way are *bona fide* persisters as they survive during antibiotic treatment, regrow after treatment, and exhibit tolerance towards antibiotics with different targets. We then used our isolation method to resolve a key outstanding question in the persistence field. Single-cell recovery of persisters after treatment was monitored in a microfluidic ‘mother machine’ device. These data show that persister awakening occurs at a constant rate, reflecting stochasticity. Our novel approach might prove useful for future single-cell studies of persistence.

## Results and discussion

### Cephalexin treatment followed by filtration enables highly efficient isolation of persister cells

Similar to the ampicillin-lysis method of Keren et al. [3], our method distinguishes persisters from normal cells based on their antibiotic tolerance, the core feature that universally characterizes all persisters and does not make any assumptions on their physiological state or underlying mechanisms. Our approach is specifically aimed at limiting the amount of cell debris in the resulting sample, as well as shortening the antibiotic exposure time. To this end, we benefit from the killing characteristics of cephalexin, a ß-lactam that does not immediately induce lysis, but first induces severe filamentation of susceptible cells before lysis is initiated (Figure 1a; Suppl. Movie 1). Cephalexin targets penicillin-binding protein (PBP) 3, also known as FtsI, a transpeptidase that is essential for peptidoglycan synthesis during cell division [12]. Drug-tolerant persisters are not affected by cephalexin and therefore do not filament in its presence, enabling their isolation from a culture by filtration (Figure 1b).

**Figure 1.**
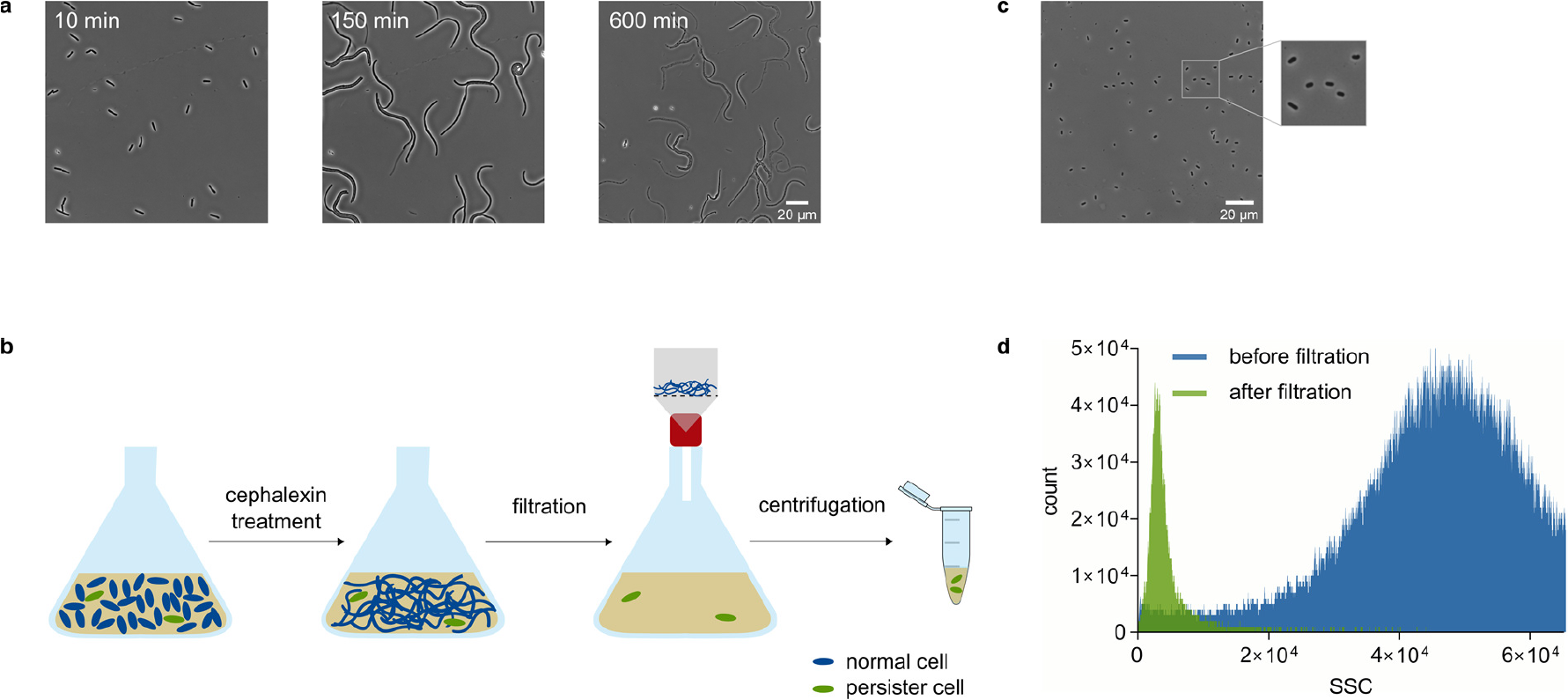
Cephalexin treatment followed by filtration enables highly efficient isolation of persister cells. (a) Susceptible, exponential phase cells filament severely during treatment with cephalexin (50 µg/ml) before lysis occurs. (b) Experimental setup of our persister isolation method. A culture in exponential phase is treated with cephalexin for one hour to induce filamentation of susceptible cells. Next, the culture is vacuum filtered (pore size of 5 µm) to separate short, antibiotic-tolerant persisters from filamented, susceptible cells. After filtration, the culture is centrifuged to remove cephalexin and to increase the density of the resulting sample. (c) Microscopic visualization of a sample after cephalexin treatment and filtration demonstrates that it mainly consists of short cells that did not respond to the cephalexin treatment. (d) Side scatter distributions of a sample before and after filtration confirm that filtration enriches for cells with the lowest side scatter values, presumably corresponding to the persisters of the culture.

ß-lactams only exhibit effective activity at low cell densities [13], implying that cultures should be in early exponential phase when cephalexin treatment starts. The biphasic killing pattern resulting from a long-term treatment with cephalexin confirms the presence of persisters in this low-density culture (Suppl. Figure S1a). By comparing the number of persisters isolated with our filtration method to the total number of persisters at the plateau of the time-kill curve (Suppl. Figure S1a), we estimated that isolation occurs with an average efficiency of 28 %. The remaining persisters are presumably lost during filtration, as filamented cells cause clogging of the filter.

Notably, the fact that filamentation occurs at a much shorter timescale than lysis considerably reduces the antibiotic exposure time as compared to the ampicillin lysis method. This was confirmed by performing our filtration protocol at different time points during a longer-term cephalexin treatment (Suppl. Figure S2a). These data show that the number of isolated cells does not change significantly for cephalexin treatments longer than one hour (p=0.17), implying that a one-hour treatment is sufficient to obtain the persisters of the culture by filtration. Any treatment shorter than one hour results in contamination with susceptible cells, while longer treatments successively generate more debris of dead cells in the sample (Suppl. Figure S2b). Indeed, an optimal treatment time of one hour results in a final sample that contains short, antibiotic-tolerant persisters and very little cell debris (Figure 1c). The purity of the resulting samples was also confirmed by the side scatter distributions of samples before and after filtration (Figure 1d).

### Cells isolated by cephalexin treatment and filtration are antibiotic-tolerant and regrow after treatment

Next, we sought to validate that cells isolated by cephalexin treatment and filtration show the key properties of persisters, being their ability to survive a longer-term antibiotic treatment and to reinitiate growth after treatment. We tested the first feature by treating a sample of isolated cells with cephalexin, both in liquid medium and on an agarose pad supplemented with rich medium (Mueller-Hinton broth; MHB). In liquid culture, cephalexin causes the number of isolated cells to decline slowly (Figure 2a), with a rate that does not significantly differ from the killing rate of persisters (p = 0.399; Suppl. Figure S1a; Suppl. Figure S1b). We hypothesize that this rate of killing, which is much lower than for susceptible cells, reflects the awakening rate of persisters in the presence of cephalexin [14]. Most cells on an MHB+agarose pad remain unaffected (Figure 2b; Suppl. Movie 2). A few isolated cells show filamentation and lysis, which can presumably be attributed to persister awakening.

**Figure 2.**
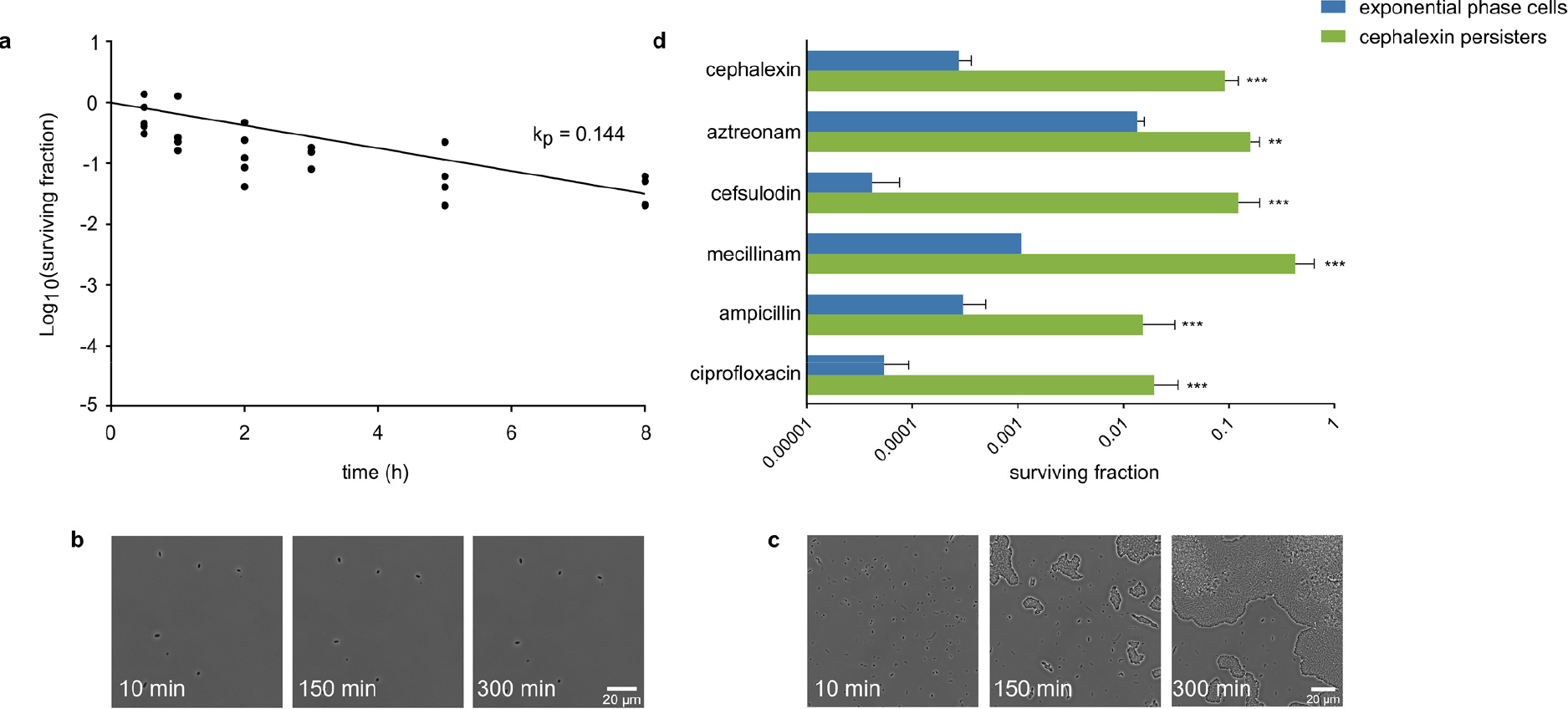
Cells isolated by cephalexin treatment and filtration are antibiotic-tolerant and regrow after treatment. (a) Time-kill curve of isolated cells treated with cephalexin (50 µg/ml) for 8 hours in liquid medium. A uniphasic exponential curve was fitted onto the data with a killing rate (*k*_*p*_ = 0.144) that is much lower than for susceptible cells (*k*_*n*_ = 4.98; Suppl. Figure S1a). The killing rate of persisters presumably reflects the rate of persister awakening in the presence of cephalexin. (b) Treatment of isolated cells with cephalexin (50 µg/ml) on an agarose pad supplemented with MHB shows that the majority of the cells is not affected by the antibiotic. (c) Persisters isolated by filtration start dividing on an agarose pad supplemented with MHB. (d) Isolated cells display multidrug tolerance, a trait associated with persistence. Fraction of surviving cells after a 5-hour treatment with cephalexin (50 µg/ml), aztreonam (0.64 µg/ml), cefsulodin (320 µg/ml), mecillinam (5 µg/ml), ampicillin (40 µg/ml), and ciprofloxacin (0.32 µg/ml), starting from an exponential phase culture or a sample consisting of isolated cephalexin persisters. For all tested antibiotics, persisters show a significantly higher tolerance as compared to exponential phase cells (cephalexin: p<0.0001, n=12; aztreonam: p=0.0004, n=9; cefsulodin: p<0.0001, n=6; mecillinam: p<0.0001, n=9; ampicillin: p<0.0001, n=9; ciprofloxacin: p<0.0001, n=9).

Importantly, these microscopic observations additionally demonstrate that the isolated cells cannot grow in the presence of cephalexin, implying that they are not genetically resistant. We then also validated that cells isolated by filtration are able to reinitiate growth, by seeding them onto an agarose pad supplemented with rich medium (Figure 2c; Suppl. Movie 3). 30-40 % of the cells started dividing within one hour, confirming their culturability after treatment. Growth of other cells was mostly masked by colonies originating from these early-dividing cells.

Persisters are often assumed to be dormant cells in which antibiotic targets are inactive, resulting in high tolerance towards various types of antibiotics. To further confirm that cells isolated by cephalexin treatment and filtration are persisters, we investigated their tolerance towards antibiotics with cellular targets that differ from PBP3. Cefsulodin is a ß-lactam that targets PBP1a and PBP1b, while mecillinam only targets PBP2. Ampicillin has multiple targets, including PBP1a, PBP1b, PBP2, and PBP3 [15]. We also investigated tolerance towards the ß-lactam aztreonam, which has the same target as cephalexin, and towards the fluoroquinolone ciprofloxacin, which targets DNA topoisomerases. Tolerance was measured by treating a sample of persisters obtained with our isolation protocol for 5 hours with the listed antibiotics. The relative fraction of surviving cells after treatment was compared to the surviving fraction of an exponential phase culture (Figure 2d). For all antibiotics, cells surviving treatment are significantly enriched in a culture that merely consists of cells isolated by filtration (334-fold for cephalexin, 12-fold for aztreonam, 2950-fold for cefsulodin, 393-fold for mecillinam, 51-fold for ampicillin, and 368-fold for ciprofloxacin). Notably, none of the tested antibiotics results in 100 % survival of the isolated cephalexin persisters. In accordance with our other data (Figure 2a), this can be partially attributed to killing of persisters as they wake up during treatment. However, these data presumably also imply that not all persisters are tolerant to all antibiotics, but rather represent a heterogeneous pool of cells with partially overlapping tolerance to various antibiotics. Together, our data show that cells isolated by cephalexin treatment and filtration are tolerant towards a longer-term cephalexin treatment, that they are able to reinitiate growth when treatment ceases, and that they show a high degree of multidrug tolerance. All these characteristics are key to the persister phenotype and make us confident that our isolated cells are *bona fide* persister cells.

### Single-cell analysis of isolated persisters in the mother machine reveals that persister awakening is a stochastic process

To microscopically examine single persister cells and their regrowth, a major drawback of using agarose pads is that early-dividing cells quickly overgrow the whole pad. First divisions of potentially later-dividing, neighbouring cells are thereby obscured, hampering quantitative single-cell analyses of awakening. To address this problem, we took advantage of the mother machine, a well-established microfluidic device that allows tracking growth of a large number of individual *E. coli* cells [16].

We isolated persisters from a culture using the filtration method described above, inserted these cells into the channels of the mother machine, and provided them with fresh nutrients (Figure 3a). Most of the channels contained either no or only one cell, allowing to track single cells. Based on a few hundred individual observations, we derived a distribution of single-cell persister awakening times for the wild-type *E. coli* strain K-12 MG1655 and the well-known high-persistence strain *hipA7* (Figure 3b). A similar distribution was obtained for both strains. This distribution shows a surprisingly high cell-to-cell variability in awakening times, ranging from a few minutes to up to 13 hours. In both cases, an exponential curve fits well to the data, indicative of a high degree of stochasticity involved in persister awakening. The persister awakening rate in fresh medium without antibiotics (*b* = 0.31-0.35; Figure 3b) is higher than in the presence of cephalexin (*k*_*p*_ = 0.04-0.18; Figure 2a; Suppl. Figure S1), although both rates were measured in different setups and therefore not perfectly comparable. While these findings corroborate existing assumptions and hypotheses about persister awakening [17,18], this is, to our knowledge, the first study that provides conclusive experimental evidence of stochastic awakening at single-cell level with such a high throughput. In addition to the awakening times, we also derived individual growth rates of freshly-awakened persisters (Suppl. Figure S3). Strikingly, these data reveal that persisters instantaneously divide at a rate that does not differ from the growth rate of normal cells. Furthermore, individual growth rates do not correlate with awakening times, indicating that cells with a long lag time do not necessarily grow slower than cells with a short lag time (Figure 3c). It should be noted that the majority of the cells did not start dividing within the course of the experiment (20 hours). As these cells are too numerous to be completely covered by the tail of the exponential distribution, they can presumably be classified as viable but non-culturable cells (VBNCs). The high abundance of VBNCs in *E. coli* cultures has already been reported before [19,20] and represents a prominent source of contamination in most persistence enrichment protocols, including ours. As VBNCs cannot be distinguished from persisters based on their antibiotic tolerance, our method is only able to discriminate between both by visualizing regrowth in fresh medium.

**Figure 3.**
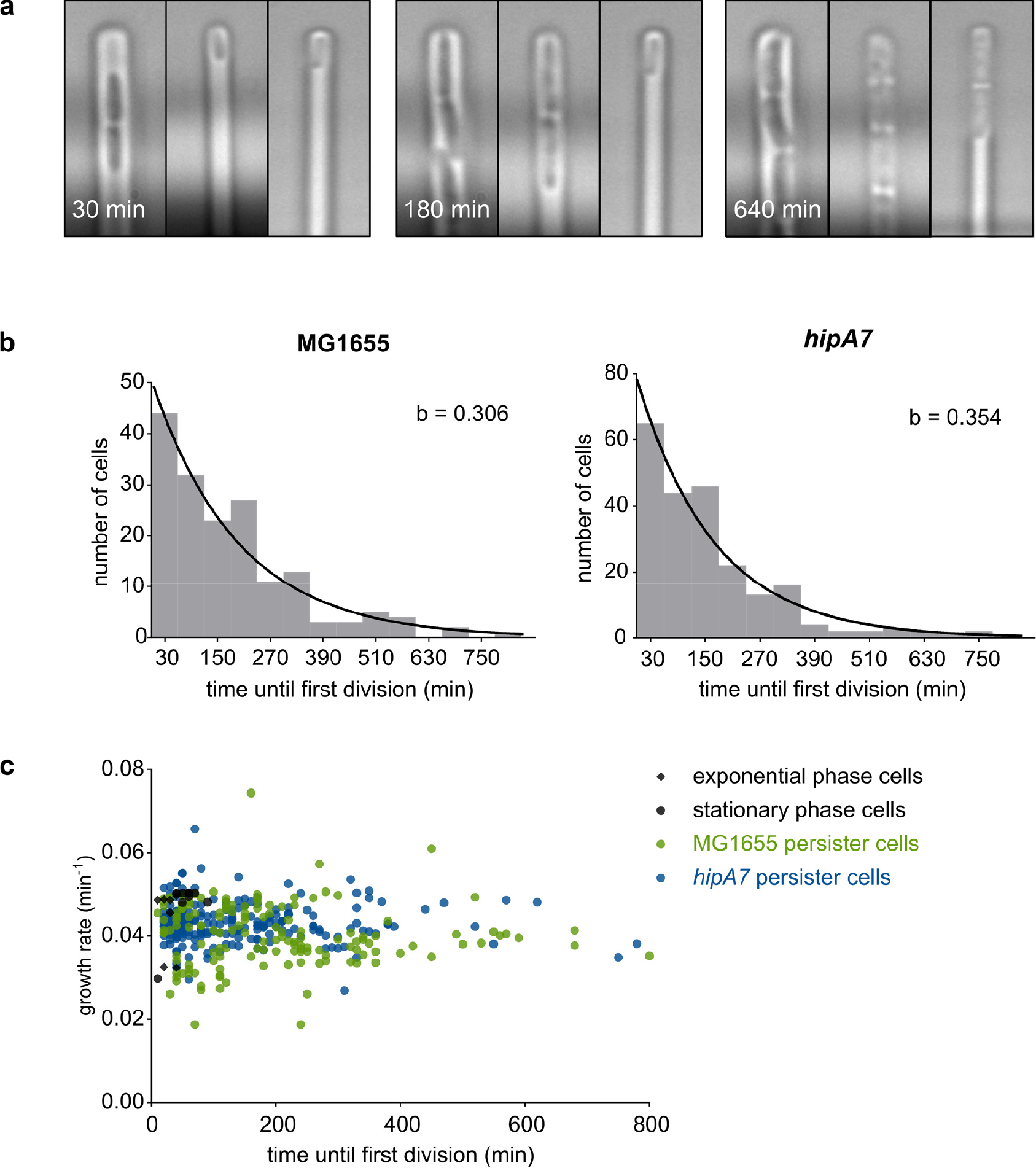
Single-cell analysis of isolated persisters in the mother machine reveals that persister awakening is a stochastic process. (a) Microscopic images of isolated persisters dividing in the mother machine. The time until first division varies strongly among individual cells (left channel: short lag time; middle channel: medium lag time; right channel: long lag time). (b) Single-cell distributions of persister awakening times measured in the mother machine, for the wild type strain MG1655 (n = 168) and the high-persistence strain *hipA7* (n = 129). An exponential distribution was fitted onto the binned data, revealing an awakening rate *b* that is similar for both strains. (c) Scatterplot of single-cell awakening times and growth rates of MG1655 and *hipA7* persisters, exponential phase cells, and stationary phase cells. While awakening times show a large variation, growth rates cluster more tightly around the average value. Both parameters are not correlated.

Cephalexin exhibits activity against a wide range of bacteria, including both Gram-positive and Gram-negative bacteria, where it elicits similar effects of filamentation and lysis. Our isolation protocol can therefore easily be extended to species other than *E. coli*. Nevertheless, the size of the inoculum used to initiate the exponential phase culture, the duration of exponential growth, and the duration and concentration of cephalexin treatment should be optimized for every strain or species, as these parameters are highly dependent on growth rate and lag phase. Ideally, an optimal balance should be found to ensure that all normal cells escaped the lag phase, while the cell density after exponential growth remains sufficiently low for an effective cephalexin treatment. We anticipate that the wide applicability of our size-separation-based persister isolation method will boost single-cell persistence studies, potentially with important consequences for the persistence field.

## Materials and Methods

### Strains, culture conditions and antibiotics

Experiments were performed with *E. coli* K-12 MG1655, except when stated otherwise. MG1655 *hipA7* was constructed by Pearl et al. [21]. Strains were grown at 37 °C in Mueller-Hinton broth (MHB) with orbital shaking (200 rpm) or on Luria-Bertani (LB) agar.

### Isolation of persisters

A 20-hour overnight culture was diluted 1:10,000 in 100 ml Mueller-Hinton broth (MHB) and incubated for 20 hours. This culture was diluted 1:5,000 in 100 ml MHB and grown for 90 minutes, to a density of 1-2 × 10^6^ CFU/ml. Next, the culture was treated with cephalexin (50 µg/ml) for 60 minutes, after which it was poured twice over a polyvinylidene fluoride membrane filter (Merck Millipore) with a pore size of 5 µm. The filtrate was collected in falcons and spun down (4,000 rpm - 5 min). After pouring off the supernatant, the remaining volume was transferred to a microcentrifuge tube and centrifuged twice (6,000 rpm - 5 min) to wash away the remaining antibiotic. The pellet was resuspended in MgSO_4_ (10 mM).

### Time-kill curves and measurement of multidrug tolerance

A 20-hour overnight culture was diluted 1:10,000 in 100 ml MHB and incubated for 20 hours. This culture was then diluted 1:5,000 in 100 ml MHB and grown for 90 minutes, to a density of 1-2 × 10^6^ CFU/ml. To measure time-kill curves of exponential phase cells, this culture was treated with cephalexin (50 µg/ml) for 16 hours. Alternatively, 1 ml of culture was transferred to a test tube and treated with cephalexin (50 µg/ml, 6× MIC), cefsulodin (320 µg/ml, 10× MIC), mecillinam (5 µg/ml, 40× MIC), aztreonam (0.64 µg/ml, 10× MIC), ampicillin (40 µg/ml, 10× MIC), or ciprofloxacin (0.32 µg/ml, 20× MIC) for 5 hours, with plating before and after treatment.

To measure time-kill curves of isolated cells, the filtration protocol was performed as stated above, after which the isolated cells were treated with cephalexin (50 µg/ml) for 8 hours. Alternatively, isolated cells were resuspended in fresh MHB and 1 ml was treated in a test tube with cephalexin (50 µg/ml, 6× MIC), cefsulodin (320 µg/ml, 10× MIC), mecillinam (5 µg/ml, 40× MIC), aztreonam (0.64 µg/ml, 10× MIC), ampicillin (40 µg/ml, 10× MIC), or ciprofloxacin (0.32 µg/ml, 20× MIC) for 5 hours, with plating before and after treatment.

### Flow cytometry

A 20-hour overnight culture was diluted 1:10,000 in 100 ml MHB and incubated for 20 hours. This culture was diluted 1:5,000 in 100 ml MHB and grown for 90 minutes, to a density of 1-2 × 10^6^ CFU/ml. Next, the culture was treated with cephalexin (50 µg/ml) for 60 minutes. A sample was taken from this culture and washed in PBS, after which the scattering values were measured by flow cytometry using a BD Influx cell sorter. The remainder of the culture was poured twice over a polyvinylidene fluoride membrane filter (Merck Millipore) with a pore size of 5 µm. The filtrate was collected in falcons and spun down (4,000 rpm - 5 min). After pouring off the supernatant, the remaining volume was transferred to a microcentrifuge tube and washed in PBS. The scattering values of this sample were measured by flow cytometry.

### Microscopy of agarose pads

To visualize killing by cephalexin, a 20-hour overnight culture was diluted 1:5,000 and incubated for 90 minutes. The resulting exponential phase culture was washed with MgSO_4_ and 2 µl of cells was seeded onto an MHB+agarose pad (2% w/v) containing cephalexin (50 µg/ml). Cells were incubated at 37 °C and killing was monitored for 6 hours. Images were obtained using a Nikon Ti-E inverted microscope with a 60× objective.

To visualize persisters, cells were isolated as described above. The resulting sample was resuspended in 10 µl MgSO_4_ and 2 µl of cells was seeded onto an MHB+agarose pad (2% w/v) with or without cephalexin (50 µg/ml). Cells were incubated at 37 °C and growth was monitored for 12 hours. Images were obtained using a Nikon Ti-E inverted microscope with a 60× objective.

### Fabrication of mother machine devices

Master molds of the microfluidic devices were designed and fabricated using standard microfabrication techniques [16]. Microfluidic chips were made by casting polydimethylsiloxane (PDMS) onto the wafer. PDMS (Sylgard 184 kit; Dow Corning) was prepared by mixing polymer base and curing agent in a 10:1 ratio. After degassing the mixture in a vacuum chamber, it was poured over the wafer and cured overnight at 65 °C. Devices were peeled from the wafer and holes for the inlet and outlet were punched using a syringe and needle (0.9 mm), and bonded to a glass coverslip after plasma activation. The bonding was established for at least 15 minutes at 65 °C. The dimensions of the growth channels were approximately 25 µm (L) × 1 µm (W) × 1 µm (D).

### Mother machine experiments

Persisters were isolated as described before and resuspended in a final volume of 20 µl MgSO_4_. After flushing the channels of the microfluidic device with MgSO_4_, cells were loaded by syringe injection followed by chip centrifugation. A peristaltic pump was used to flow medium through the device at a flow rate of 90 µl/min. The microscope chamber, which also contained the medium reservoir, was constantly held at 37 °C. Images were obtained using an Olympus XI71 inverted microscope with a 100× objective.

### Data analysis and statistics

Images were analysed using NIS Elements D 4.60.00 (Nikon Instruments, Japan) and ImageJ (https://imagej.nih.gov/). Flow cytometry data were analysed with FlowJo V10. Statistical analyses were performed in R (https://www.r-project.org/).

#### Time-kill curves

Biphasic killing parameters were determined by fitting a bi-exponential mixed model to the Log_10_-transformed, normally distributed number of surviving cells (CFU/ml) using the R package *nlme* (https://cran.r-project.org/web/packages/nlme/index.html). The model was based on the equation Log_10_(CFU)=Log_10_(N_0_.e^−kn.τ^ + P_0_.e^−kp.τ^), with τ the treatment time (in hours), N_0_ and P_0_ the number of normal and persister cells at τ=0, and k_n_ and k_p_ the rate at which normal and persister cells are killed (per hour) [22]. For uniphasic killing, the R package *lme4* (https://cran.r-project.org/web/packages/lme4/index.html) was used to fit the equation Log_10_(CFU)=Log_10_((N_0_+P_0_).e^−k.τ^). AIC (Akaike Information Criterion) was used to assess both models.

#### Multidrug tolerance

Surviving fractions were compared between conditions using one-way ANOVA and post-hoc comparisons with Sidak’s correction for multiple testing.

#### Distribution of awakening times

The variable ‘persister awakening time’ was split into bins of 60 minutes. The number of observations in each bin was normalized, to obtain relative frequencies (*freq*) of awakening events. The *nls* function in R was used to fit an exponential distribution with equation Log_10_(*freq*) = Log_10_(b.e^−b.τ^) onto the data, with τ the time in fresh medium and b the rate of awakening. After checking normality with a Shapiro-Wilk test, Log_10_-transformed awakening times were compared statistically among different strains or cell types using an unpaired, two-sided *t*-test, with Welch correction in the case of unequal variances (checked with an F-test).

#### Growth rates

A piecewise linear function was fitted to the cumulative number of divisions over time, for each individual cell. The number of knots was chosen by cross-validation and most often corresponds to one, dividing the growth curve into a lag phase and exponential growth phase. The slope of the second curve was then used to derive the average growth rate of individual cells. After checking normality with a Shapiro-Wilk test, growth rates were compared statistically among different strains using an unpaired, two-sided *t*-test, with Welch correction in the case of unequal variances (checked with an F-test).

## Data availability

The data that support the findings of this study are available from the corresponding author upon request.

## Acknowledgements

We thank Abram Aertsen (KU Leuven) for sharing the *hipA*7 strain. E.M.W. and B.V.d.B are Research Foundation Flanders (FWO)-fellows. This research was funded by the KU Leuven Research Council (PF/10/010; C16/17/006), FWO (G047112N; G0B2515N; G055517N), and the Flemish Institute for Biotechnology (VIB).

## Author contributions

E.M.W. designed and performed the experiments, analyzed the data and wrote the manuscript. Z.B.M helped performing the experiments. T.Z., B.V.d.B., P.H. and J.M. helped designing the experiments and edited the manuscript. K.J.V. edited the manuscript.

---

**Figure S1.**
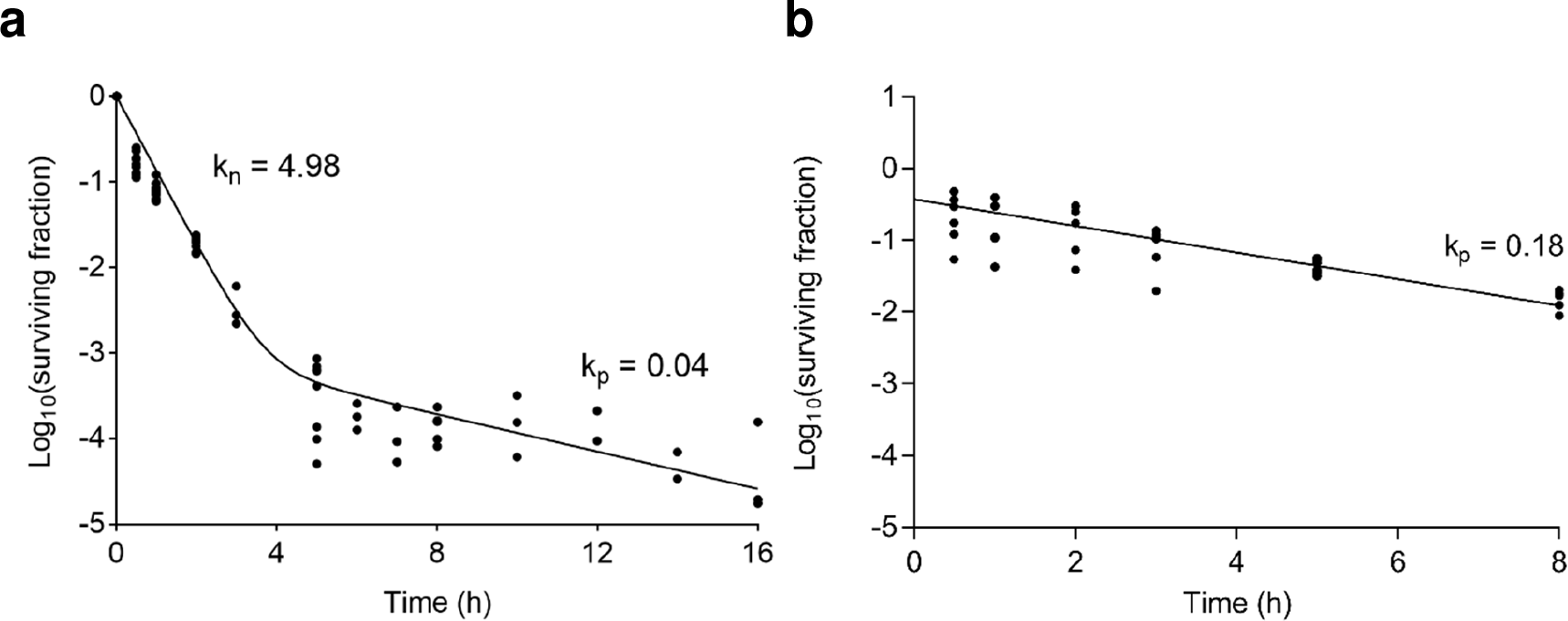
Persisters are killed slowly in the presence of cephalexin. (a) Time-kill kinetics of an exponential phase culture treated with cephalexin (50 µg/ml) for 16 hours. A biphasic exponential curve was fitted onto the data, with the first phase representing the fast killing rate of susceptible cells (*k*_*n*_), and the second phase showing the slow killing rate of tolerant persisters (*k*_*p*_). (b) A culture was first treated with cephalexin (50 µg/ml) for 5 hours to kill all susceptible cells (not shown). The remaining persisters were then exposed to an 8-hour cephalexin treatment (50 µg/ml). A uniphasic exponential curve was fitted onto the data, with the killing rate (*k*_*p*_) presumably representing the awakening rate of persisters. This rate does not differ significantly from the killing rate of cells isolated by cephalexin treatment and filtration (p = 0.399; Figure 2a of the main text). Best-fit estimated values of the killing parameters are indicated on the graphs.

**Figure S2.**
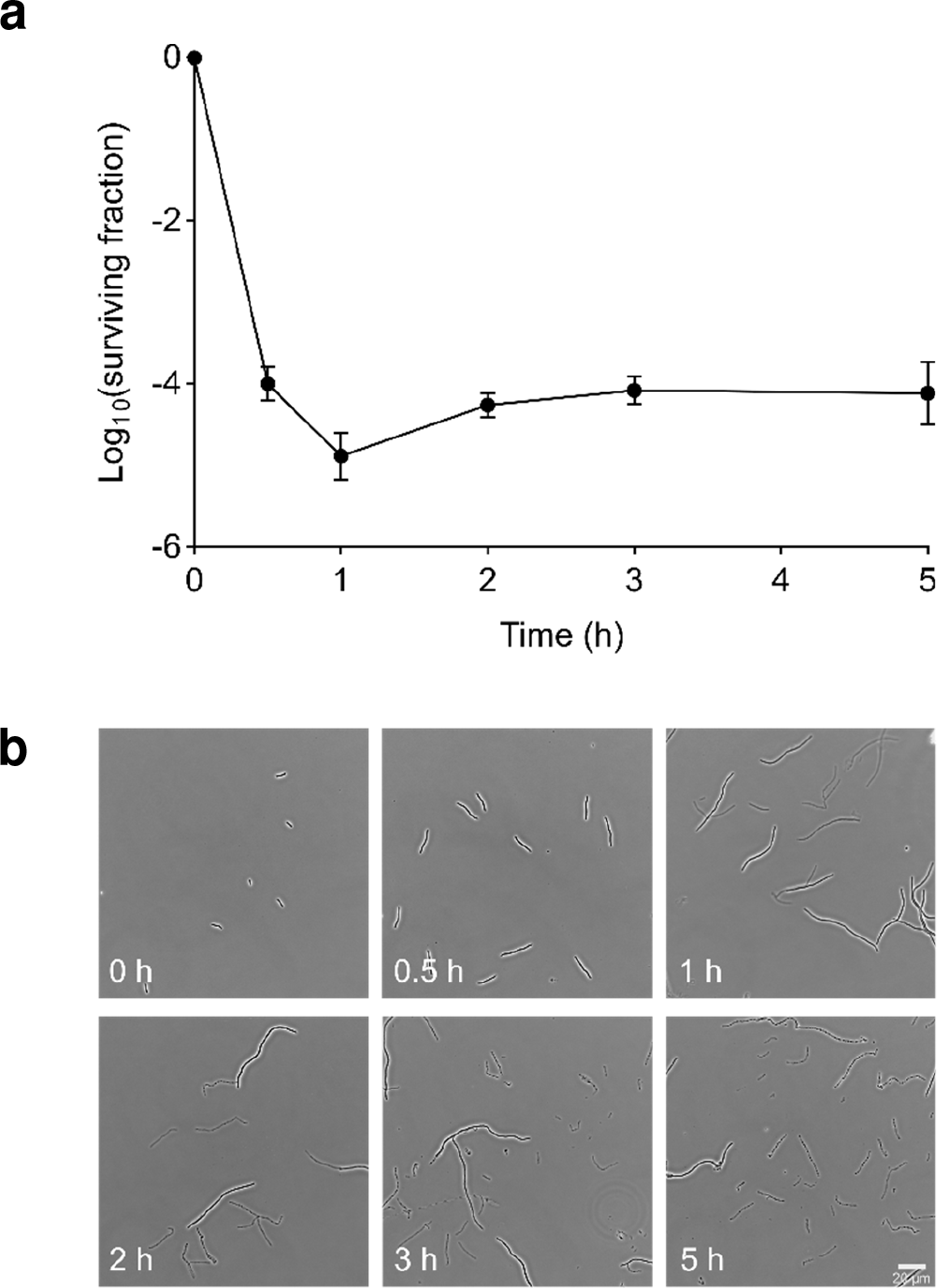
A 1-hour cephalexin treatment is optimal for persister isolation by filtration. (a) Filtration was performed at regular time points during a 5-hour cephalexin treatment (50 µg/ml). After 30 minutes of treatment, the sample after filtration is still contaminated with susceptible cells due to insufficient filamentation. A treatment of 1 hour is sufficient to obtain a sample only containing persisters, as the number of cells does not decrease further when the treatment is extended (n=3). The latter was confirmed by fitting a linear model to the data (time ≥ 1 h). The slope of this model is not significantly different from zero (p = 0.17). (b) Microscopy images of samples taken at different time points during treatment of an exponential phase culture with cephalexin (50 µg/ml), without performing filtration. A treatment longer than 1 hour results in an increasing amount of debris from lysed cells and might therefore hamper subsequent single-cell studies.

**Figure S3.**
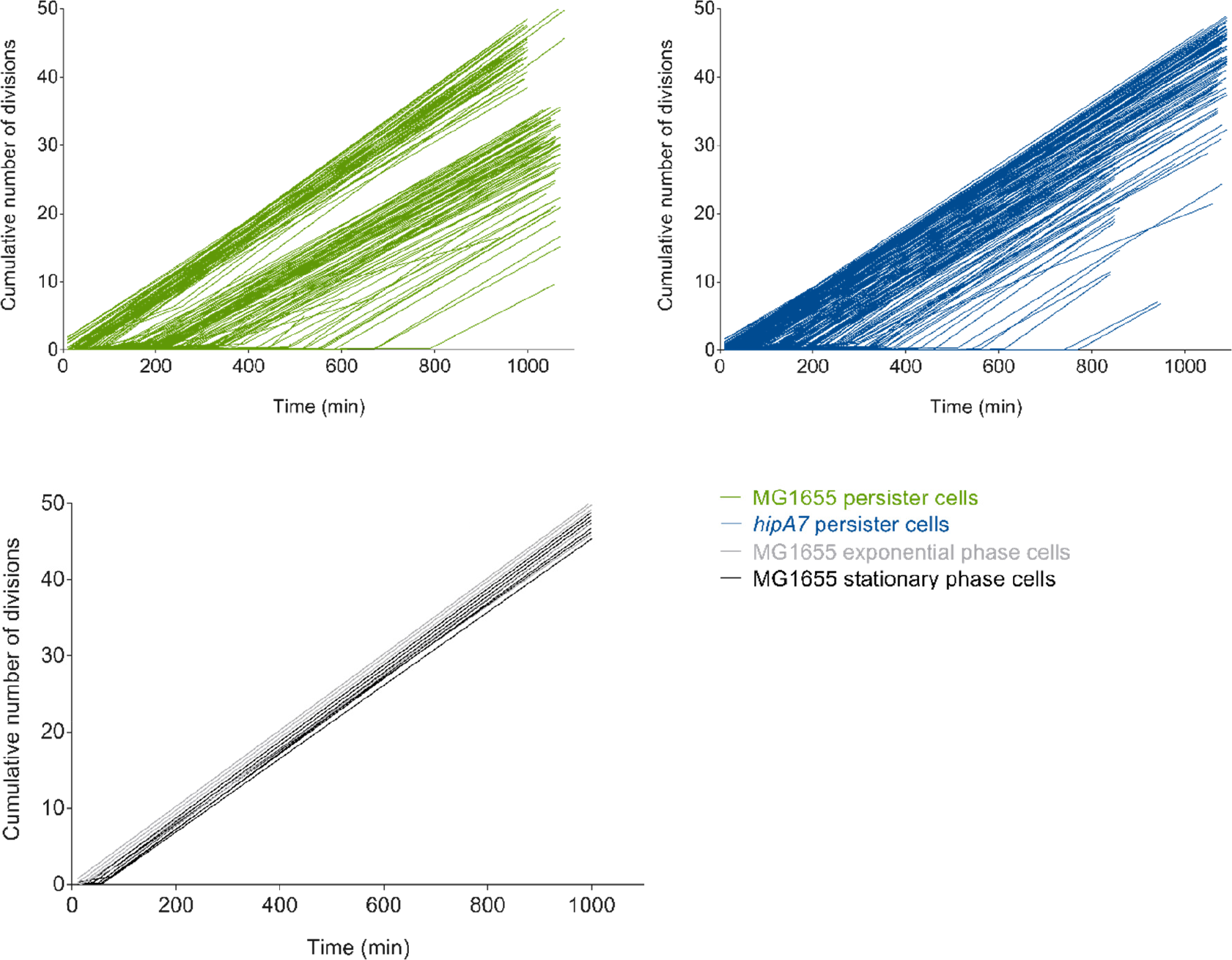
Fittings to the cumulative number of divisions of awakened persisters in the mother machine. Linear splines were fitted onto the cumulative number of divisions for each awakened persister, as well as for exponential and stationary phase cells observed in the mother machine. Individual growth rates were derived from the slopes of the fitted curves (MG1655 persisters: n=168; *hipA7* persisters: n=129; MG1655 exponential phase cells: n=10; MG1655 stationary phase cells: n=11).

